# A data citation roadmap for scientific publishers

**DOI:** 10.1101/100784

**Authors:** Helena Cousijn, Amye Kenall, Emma Ganley, Melissa Harrison, David Kernohan, Thomas Lemberger, Fiona Murphy, Patrick Polischuk, Simone Taylor, Maryann Martone, Tim Clark

## Abstract

This article presents a practical roadmap for scholarly publishers to implement data citation in accordance with the Joint Declaration of Data Citation Principles (JDDCP), a synopsis and harmonization of the recommendations of major science policy bodies. It was developed by the Publishers Early Adopters Expert Group as part of the Data Citation Implementation Pilot (DCIP) project, an initiative of FORCE11.org and the NIH BioCADDIE program. The structure of the roadmap presented here follows the “life of a paper” workflow and includes the categories Pre-submission, Submission, Production, and Publication. The roadmap is intended to be publisher-agnostic so that all publishers can use this as a starting point when implementing JDDCP-compliant data citation. Authors reading this roadmap will also better know what to expect from publishers and how to enable their own data citations to gain maximum impact, as well as complying with what will become increasingly common funder mandates on data transparency.

## Introduction

Over the past several years many authoritative science policy bodies have recommended robust archiving and citation of primary research data to resolve problems in reproducibility, robustness and reusability. Studies by CODATA (https://www.codata.org), the U.S. National Academy of Sciences, the Royal Society, and other groups recommend that scholarly articles now treat the primary data upon which they rely as first class research objects^1^-^5^. Primary data should be robustly archived and directly cited as support for findings, just as literature is cited; and where data is re-used for subsequent analysis, it should be cited as well, thus recognising the value of the data, and ensuring credit to those who generated the data. The archived data is strongly recommended – as a matter of good scientific practice - to be “FAIR”: Findable, Accessible, Interoperable, and Reusable; and to be accessible from the primary article^6^. A widely recommended method for establishing this accessibility is by data citation.

The Joint Declaration of Data Citation Principles (JDDCP) summarizes the recommendations of these studies, and has been endorsed by over 100 scholarly organizations, funders and publishers^7^. Further elaboration on how to implement the JDDCP was provided in Starr et al. 2015^8^, with an emphasis on accessibility practices for digital repositories. There is a clear emerging consensus in the scholarly community, including researchers, funders, and publishers, supporting the practice of data archiving and citation. This is reflected not only in the broad endorsement of the JDDCP, but also in the increasing proliferation of workshops on this topic. At least one journal, which had earlier published a widely discussed editorial by clinical trialists arguing against openly sharing data, is now leading an effort to help provide institutional incentives for authors to share and cite data^9^.

There is also evidence to suggest that researchers, primarily to enhance the visibility and impact of their work, but also to facilitate transparency and encourage re-use, are increasingly sharing their own data, and are making use of shared data from other researchers^10^. Researchers with funding that requires open data therefore need to know what to expect from publishers that support data citation. Which databases or repositories are acceptable places to archive their data? Should they deposit in institutional repositories, general-purpose repositories such as Dataverse, Dryad, or Figshare, or domain-specific repositories? How are embargoes handled? Can confidential data or data requiring special license agreements for sharing, be archived and cited? How should the citation itself be formatted? We attempt to answer these and other key questions in this article.

While intellectual property and confidentiality remain important considerations for researchers as potential inhibitors to sharing, researchers are also concerned about receiving appropriate citation credit or attribution for major data production efforts. We hope to provide a standardized route to clear and accessible data citation practices, which should help to alleviate most authors’ concerns and clear up any potential confusion about sharing.

The growing expectation of authors to support the assertions made in their research articles in order to maintain transparency and reproducibility requires depositing the underlying data in openly accessible locations (which are not Supplementary files to the research article). There is evidence to support this as a benefit to authors by increasing citations and usage ^11,12^ as well as scientific progress itself ^13^. The additional benefit to authors is that their data are more likely to sit alongside appropriate material in a site-specific repository, as well as receiving guidance from the repository managers and curators. Many databases are well equipped to ensure publication of datasets is timed with publication of the associated research article.

Funders and research institutions increasingly will require full primary data archiving and citation. Publishers must therefore adapt their workflows to enable data citation practices and provide tools and guidelines that improve the implementation process for authors and editors, and relieve stress points around compliance. One approach that has been taken as a means to recognising data as a first-class research object, is to create Data journals, such as Nature Scientific Data, Data in Brief, and Gigascience. These journals oversee peer review of a publication about the dataset itself and its generation; publication of a data descriptor paper results. However, this article presents a path for other journals to implement data citation developed by a team of experts from leading early adopters of data citation in the publishing world who have collectively outlined a standard model. It covers all phases of the publishing life cycle, from instructions to authors, through internal workflows and peer reviewing, down to digital and print presentation of content.

Implementing data citation is not meant to replace or bypass citation of the relevant literature, but rather to ensure we provide verifiable and re-usable data that supports published conclusions and assertions. Data citation is aimed at significantly improving the robustness and reproducibility of science; and enabling FAIR data at the point of its production. The present document is a detailed roadmap to implementing JDDCP-compliant data citation, prepared by publishers, for an audience of publishers and authors, as part of a larger effort involving roadmap and specification development for and by repositories, informaticians, and identifier / metadata registries^14,15^. We hope, in the long run, that open data will become a common enough practice so that all authors will eventually expect to provide it and cite it, and that this practice will be supported by all publishers as a matter of course.

## Results

This section briefly explains data citations and presents implementation recommendations for publishers, editors and scholarly societies. Although throughout this roadmap we refer to implementation falling under the remit of the publisher, due to the diversity of publishing models, this might not always be the case. Where an aspect of implementation falls to another party (e.g., a society journal where journal policy would often be set by the society), approval of and participation in implementation from that party would be needed.

Data citations are formal ways to ground the research findings in a manuscript upon their supporting evidence, when that evidence consists of externally archived datasets. They presume that the underlying data has been robustly archived in an appropriate long-term- persistent repository. This approach supersedes “Supplemental Data” as a home for article- associated datasets. It is designed to make data fully FAIR (Findable, Accessible, Interoperable and Reusable).

Publishers implementing data citation will provide domain-specific lists of acceptable repositories for this purpose, or guide authors to sites that maintain these lists. We provide examples of some of these lists further along in the manuscript. Guidance on why and how to cite data for authors can be found in Table 1. Formatting guidance will differ by publisher and by journals, but some examples of data citation reference styles can be found in Box 1. Figure 1 illustrates a data citation. Figure 2 shows the ideal resolution structure from data citations, to dataset landing pages, and to archived data.

**Table 1.** How and why to cite data

| **Why and How to Cite Data – For Authors** |  |
| --- | --- |
| *Why?* | 1. Supports reproducibility and validation of results, allows data reuse.<br>2. Provides credit for data generators.<br>3. Publications linked to publicly available data have been associated with increased citations.<br>4. Improves connectivity and provenance tracking of data described in publications.<br>5. Data sharing is increasingly required by funders, publishers, and institutions. |
| *How?* | 1. For primary data: determine an appropriate long-term repository for data archiving. Your publisher should provide access to a list of acceptable archival repositories.<br>2. Deposit your data and get an accession number or dataset DOI from the repository.<br>3. For secondary data: cite what you use. When using the data of others, cite both the related peer-reviewed literature and the actual datasets used.<br>4. Include ‘formal’ data citations whenever possible: When datasets have formal, stable identifiers or accession numbers, they should be included in the main reference list.<br>5. Be as complete as possible, but don’t invent metadata: If a data record does not have a clear author/creator or title, don’t make one up.<br>6. Refer to your publisher’s *Guide for Authors* to format your dataset reference. |
| **Why and How to Cite Data – For Publishers** |  |
| *Why?* | 1. Help authors and journals to comply easily with funder mandates.<br>2. Improve author service by simplifying policies and procedures and increasing the visibility and connectivity of their articles and data.<br>3. Improve editor and peer reviewer service with better guidelines and support for data and visibility of data in the peer review process.<br>4. Improve reader and author service with more consistent links to data.<br>5. Support editorial goals to publish more open and reproducible research.<br>6. Make the most of your repository partnerships. |
| *How?* | 1. Revise editor training and advocacy material<br>2. Revise reviewer training material<br>3. Update information for authors by:<br>a. providing guidance on author responsibilities and a policy on data citation;<br>b. asking authors to provide a Data Availability Statement;<br>c. specifying how to format data citations; and<br>d. providing detailed guidance on suitable repositories.<br>4. Update guidelines for internal customer services queries and provide author FAQs.<br>5. Capture data citation in reference list at point of submission in a structured way.<br>6. Data availability should be captured in a structured way.<br>7. Update XML DTD for data citation tagging.<br>8. Display data citations in the article.<br>9. Deliver data citation metadata to Cross-Ref. |

**Figure 1.**
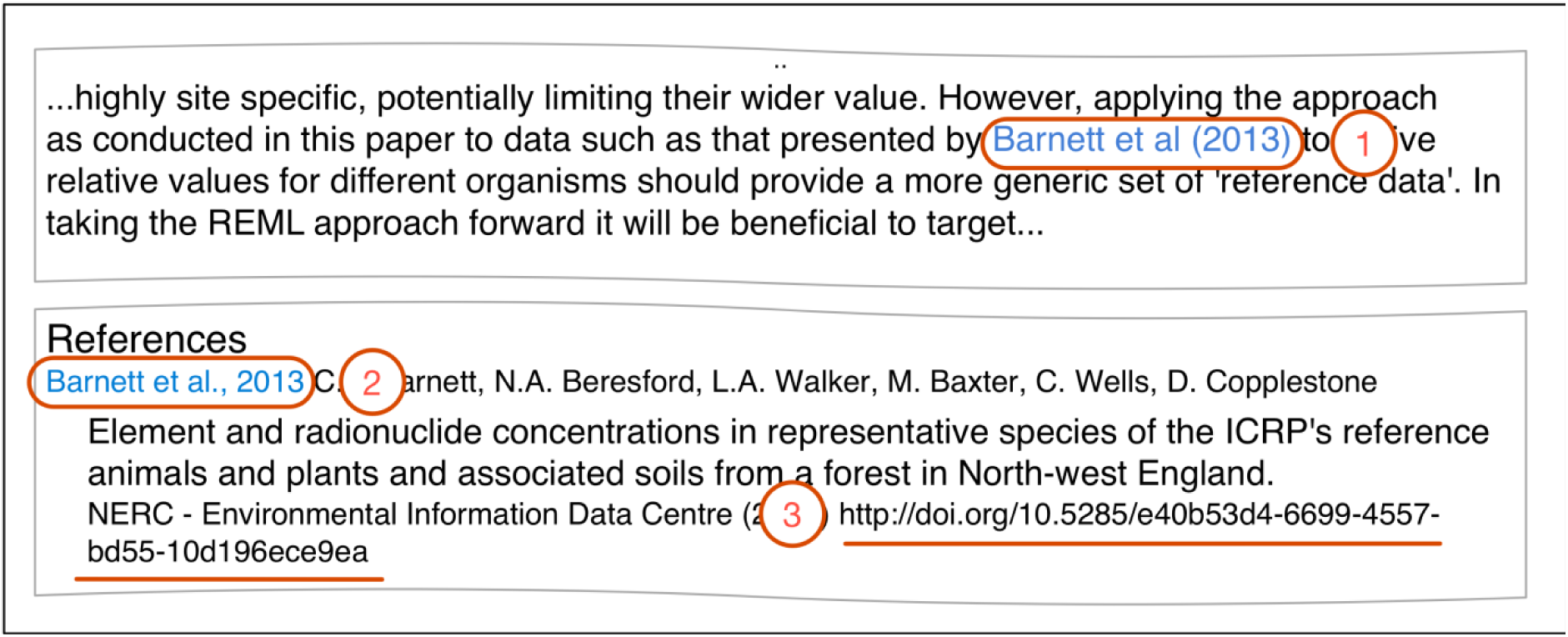
Data citation example. (1) Data citation in text; (2) Reference; (3) Globally resolvable unique identifier. *Example from Beresford NA, et al. (2016) Journal of Environmental Radioactivity 151(2): 373-386. https://doi.org/10.1016/j.jenvrad.2015.03.022.*

**Figure 2.**
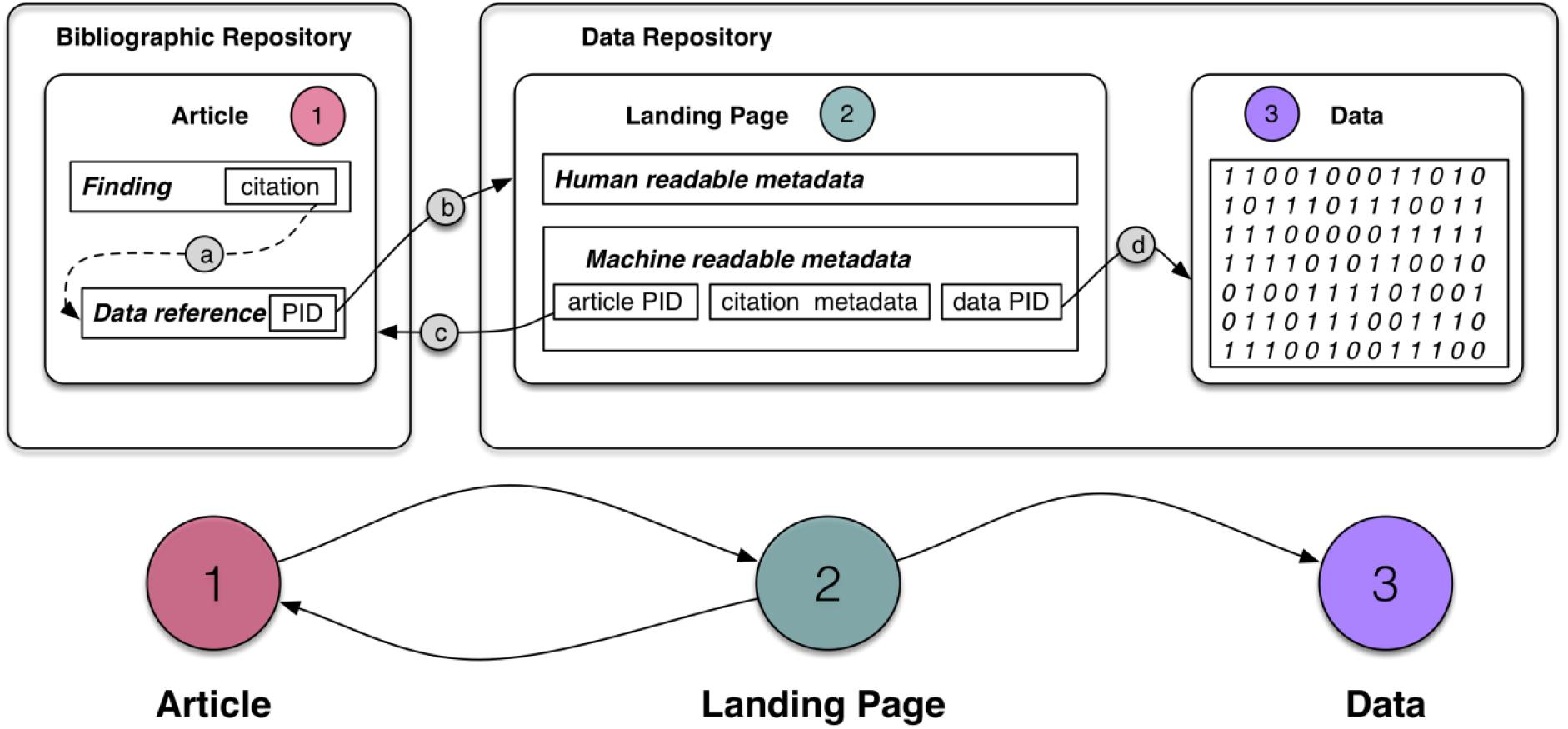
Data citation resolution structure (ideal workflow). Articles (1) link to datasets in appropriate repositories, on which their conclusions are based, through citation to a dataset (a), whose unique persistent identifier (PID) resolves (b) to a landing page (2) in a well-supported data repository. The data landing page contains human- and machine-readable metadata, to support search and to resolve (c) back to the citing article, and (d) a link to the data itself (3).

#### Box 1. Reference style examples for citing data

Numbered style:

[dataset] [27] M. Oguro, S. Imahiro, S. Saito, T. Nakashizuka, Mortality data for Japanese oak wilt disease and surrounding forest compositions, Mendeley Data, v1, 2015. https://doi.org/10.17632/xwj98nb39r.1

[dataset] [28] D. Deng, C. Xu, P.C. Sun, J.P. Wu, C.Y. Yan, M.X. Hu, N. Yan, Crystal structure of the human glucose transporter GLUT1, Protein Data Bank, 21 May 2014. https://identifiers.org/pdb:4pyp

Harvard style:

[dataset] Farhi, E., Maggiori, M., 2017. “Replication Data for: ‘A Model of the International Monetary System’”, Harvard Dataverse, V1. https://doi.org/10.7910/DVN/8YZT9K

[dataset] Aaboud, M, Aad, G, Abbott, B, Abdallah, J, Abdinov, O, Abeloos, B, AbouZeid, O, Abraham, N, Abramowicz, H, Abreu, H., 2017. Dilepton invariant mass distribution in SRZ. HEPData, 2017-02-08. https://doi.org/10.17182/hepdata.76903.v1/t1

Vancouver style:

[dataset] [52] Wang G, Zhu Z, Cui S, Wang J. Data from: Glucocorticoid induces incoordination between glutamatergic and GABAergic neurons in the amygdala. Dryad Digital Repository, August 11, 2017. https://doi.org/10.5061/dryad.k9q7h

[dataset] [17] Polito VA, Li H, Martini-Stoica H, Wang B et al. Transcription factor EB overexpression effect on brain hippocampus with an accumulation of mutant tau deposits. Gene Expression Omnibus, December 19, 2013. https://identifiers.org/GEO:GDS5303

APA style:

[dataset] Golino, H., Gomes, C. (2013). Data from the BAFACALO project: The Brazilian Intelligence Battery based on two state-of-the-art models: Carroll’s model and the CHC model. Harvard Dataverse, V1, https://doi.org/10.7910/DVN/23150

[dataset] Justice, L. (2017). Sit Together and Read in Early Childhood Special Education Classrooms in Ohio (2008-2012). ICPSR 36738.

https://doi.org/10.3886/ICPSR36738.v1

AMA style:

[dataset] 12. Kory Westlund, J. Measuring children’s long-term relationships with social robots. Figshare, v2; 2017. https://doi.org/10.6084/m9.figshare.5047657

[dataset] 34. Frazier, JA, Hodge, SM, Breeze, JL, Giuliano, AJ, Terry, JE, Moore, CM, Makris, N. CANDI Share Schizophrenia Bulletin 2008 data; 2008. Child and Adolescent NeuroDevelopment Initiative. https://dx.doi.org/10.18116/C6159Z

Both the dataset reference in the primary article, including its globally resolvable unique persistent identifier (PID), and the archival repository, should follow certain conventions. These are ultimately based upon the JDDCP’s eight principles. Initial conventions for repositories were developed in *Starr et al 2015*^8^ and are presented in more depth and detail in *Fenner et al. 2017* ^15^.

The remainder of this article is organized as a set of proposed actions for publishers, and linked to author responsibilities, applicable to each point in the lifecycle of a research article: Pre-submission, Submission, Production, and Publication.

### 1. Pre-submission

#### 1.1 Revise editor training and advocacy material

Editor advocacy and training material should be revised. This may differ by journal or discipline, and whether there are in-house editors, academic editors, or both. For example, this might involve updates to the editor training material (internally maintained, for example, on PowerPoint or PDFs, or externally on public websites) or updates to advocacy material. The appropriate material should be revised to enable editors to know what data citation is, why it should be done, what data to cite, and how to cite data. This should equip editors to instruct reviewers and authors on journal policy around data citation.

#### 1.2 Revise reviewer training material

Reviewer training material should be revised to equip reviewers with the knowledge needed to know what data authors should cite in the manuscript, how to cite this data and how to access the underlying data to a manuscript. Training material should also communicate expectations around data review. Several other projects are underway focusing on defining criteria for data peer review.

#### 1.3 Update information for authors

##### Provide guidance on author responsibilities

Data citation is based on the idea that the data underlying scientific findings or assertions should be treated as first-class research objects. This begins with author responsibility to properly manage their own data prior to submission. The Corresponding Author should have ultimate oversight responsibility to ensure this is done in a transparent, robust and effective way ^16^. Researchers are also increasingly required by funders to submit data management plans. No later than the time of submission (and ideally at the time of data generation), researchers should take responsibility for determining an appropriate repository that supports data citation (with landing pages, PID, and versioning) and provides support to ensure appropriate metadata are present. The publisher’s responsibility in this regard, is to provide or refer authors to a definitive list of such repositories in the *Guide for Authors*.

##### Specify a policy for data citation

Data citation should be implemented at a journal policy level, as part of a journal’s wider policy on data sharing. It is recommended that this policy, since it is discipline-specific, should be determined by the journal community (editor, reviewers, etc.) as well as the publisher.

There are multiple options for a data policy. For example, Springer Nature, Wiley and Elsevier have all recently rolled out a range of multi-level policies depending on their specific needs ^17^-^19^. This means that they offer their journals a range of policy options ranging from encouragement of data sharing, to strong encouragement, to mandatory data sharing. Additionally, data policies can also be defined at the domain level as was done by COPDESS (the Coalition for Publishing Data in the Earth & Space Sciences), an initiative within the geosciences ^20^. Another approach, taken by the Public Library of Science (PLOS), was to have a single policy requiring that all underlying data be made available at the time of publication with rare exception for all of their journals ^21^. Whatever the level of the policy, it should specify which datasets to cite (e.g., underlying data versus relevant data not used for analysis) and how to format data citations. Authors should provide details of previously published major datasets used and also major datasets generated by the work of the paper. It is recommended if at all possible that data citation occurs either in the standard reference list or (less preferable) in a separate list of cited data, formatted similarly to standard literature references. But regardless of where citations appear in the manuscript, they should be in readily parsable form and therefore machine readable.

##### Ask authors for a Data Availability Statement (DAS)

It is recommended that as part of data citation implementation publishers adopt standardized Data Availability Statements (Figure 3). DASs provide a statement about where data supporting the results reported in a published article can be found, including, where applicable, unique identifiers linking to publicly archived datasets analyzed or generated during the study. In addition, DASs can increase transparency by providing a reason why data cannot be made immediately available (such as the need for registration, due to ethical or legal restrictions, or because of an embargo period). Some research funders, including Research Councils UK, require data availability statements to be included in publications so it is an important element of a publisher’s data policy. It is recommended that publicly available datasets referred to in DASs are also cited in reference lists.

**Figure 3.**
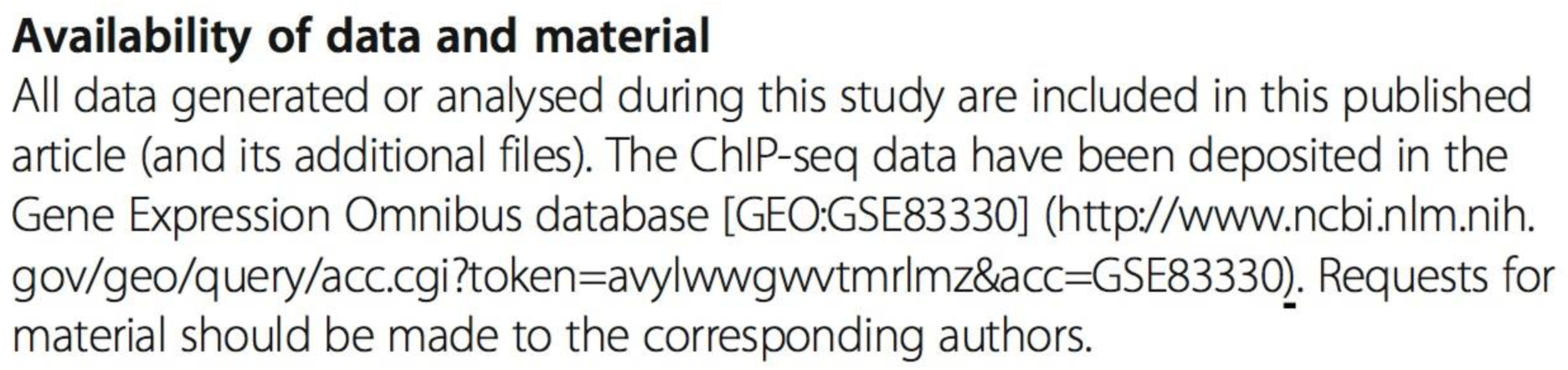
Example of a Data Availability Statement. Taken from Kim et al. 2016^22^. Available at https://doi.org/10.1186/s12915-016-0320-z.

##### Specify how to format data citations

Whilst there are many referencing style guides, including formal standards managed by ISO/BS (<u>ISO 690-2010</u>) and ANSI/NISO (<u>NISO Z39.29-2005 R2010</u>), several of the key style guides provide guidance on how to cite datasets in the reference list. In addition, the reference should also include the tag “[dataset]” within the reference citation so that it becomes easily recognizable within the production process. This additional tag does not have to be visible within the reference list of the article. It is critical to ensure the recommended format of the data citation also adheres to the Joint Declaration of Data Citation Principles. Publishers should provide an example of the in-text citation and of the reference to a dataset in their references formatting section (see Box 1 for examples).

Similar to article references, key elements for data citation include, but may not be limited to: author(s), title, year, version, data repository, PID. Further information can be found on the DataCite website (https://www.datacite.org/cite-your-data.html). Researchers should refer to journal-specific information for authors on publisher websites for definitive guidance on how to cite data when submitting their manuscript for publication.

##### Provide guidance around suitable repositories (general, institutional, and subject-specific) and how to find one

Publishers should provide or point to a list of recommended repositories for data sharing. Many publishers already maintain such a list. The Registry of Research Data Repositories (Re3Data, https://www.re3data.org) is a full-scale resource of registered repositories across subject areas. Re3Data provides information on an array of criteria to help researchers identify the ones most suitable for their needs (licensing, certificates & standards, policy, etc.). A list of recommended repositories is provided by FAIRsharing.org, where some publishers also maintain collections of recommended resources. FAIRsharing started out as a resource within the life sciences but has recently expanded, and now includes repositories within all disciplines.

Where a suitable repository does not exist for a given discipline or subject area, publishers should provide guidance for the use of a general purpose or institutional repository where these meet the recommendations of the repository draft roadmap guidance ^15^ (briefly, by providing authors’ datasets with a globally resolvable unique identifier - ideally a DataCite DOI where possible, or other PID, providing a suitable landing page, using open licenses, and ensuring longevity of the resource).

Some research funders may stipulate that data must be deposited in a domain-specific repository where possible, which aligns well with publishers providing lists of recommended repositories.

Examples of publisher- or consortium-maintained recommended repositories lists include:

- PLOS: http://journals.plos.org/plosbiology/s/data-availability#loc-recommended-repositories
- SpringerNature: http://www.springernature.com/gp/group/data-policy/repositories
- EMBO Press: http://emboj.embopress.org/authorguide#datadeposition
- Elsevier: https://www.elsevier.com/authors/author-services/research-data/data-base-linking/supported-data-repositories
- COPDESS: https://copdessdirectory.osf.io

Fairsharing.org is currently working with several publishers to develop and host a common list, which is expected to be available later in 2018. At that time, participating publishers hope to link directly to the single recommended list from their author instructions.

##### Provide specific guidance on in-text accessions or other identifiers, particularly in citing groups of datasets reused in meta-analyses

Publishers should provide guidance to authors on dealing with list of accessions or other identifiers in text. This is especially relevant for re-used datasets in meta-analysis studies. Particularly, in the biomedical sciences, meta-analyses may reuse a large number of datasets from archives such as the Gene Expression Omnibus (GEO)^23,24^. When many input datasets need to be cited, authors should use the EMBL-EBI’s Biostudies database^25^ or similar, to group the input accessions under a single master accession for the meta-analysis, and they should then cite the master accession for the Biostudies entry in their reference list. Supplements or Appendices should not be employed for this purpose.

In general, any lists of accessions or other identifiers appearing in the text should be accompanied by appropriate data citations, mapped to appropriate entries in the Reference list, or grouped in a Biostudies or similar entry and cited as a group. Editors should receive guidance from the Publisher on how to promote this approach.

##### Consider licensing included under “publicly accessible” and implications (e.g. automated reuse of data)

Publishers should consider the types of licensing allowed under their data policy. It is recommended that data submitted to repositories with stated licensing policies should have licensing that allows for the free reuse of that data, where this does not violate protection of human subjects or other overriding subject privacy concerns. Many publishers use Creative Commons licenses as a guide for equivalence criteria that repository licenses should meet.

##### Update guidelines for internal customer services queries and provide author FAQs

Publishers will need to include a support service around their data policy. This might include a list of author-focused FAQs. Internal FAQs should also be provided to customer services. Alternatively, or in addition, publishers might set up a specific email address for queries concerning data. PLOS, Springer Nature and Elsevier provide such email addresses.

### 2. Submission and review

#### Cite datasets in text of manuscript, and present full data citations in the reference list

At the submission stage it is important that all the required elements are captured to create a data citation: author(s), title, year, version, data repository, PID. The recommended way to capture data citations is to have authors include these in the reference list of the manuscript. Instructions for data citation formatting can be found in the pre-submission section above and will depend on the reference style of the journal. In all cases, datasets should be cited in the text of the manuscript and the reference should appear in the reference list. To ensure data references are recognized, authors should indicate with the addition of “[dataset]” that this is a data reference (see examples in **Box 1** above).

#### Data availability should be captured in a structured way

At the time of submission, authors should be requested to include a DAS about the availability of their data. In situations where data cannot be made publicly available, this should be explained here. This statement can be used to detail any other relevant data-related information. The JATS for Reuse (JATS4R) group has produced a draft recommendation for tagging data availability statements (http://jats4r.org/data-availability-statements). This group recommends the statement is separate and not displayed as part of the acknowledgements.

#### Editors and reviewers are enabled to check the data citation and underlying data

Through the data citation, editors and reviewers should be able to access underlying datasets. Datasets on which any claims in an article are based, should always be available to peer reviewers. If researchers do not want their data to be public ahead of the manuscript’s publication, some repositories can provide a reviewer access link. If available, this should be provided at the time of submission. If the data are not available from the repository during review, authors should be willing to work with the Publisher to provide access in another mutually agreeable manner. Reviewer forms should be updated with information on how to access the data and a question about whether data sharing standards/policies have been met. Publishers should be mindful that they do not reveal the identity of the authors in cases where peer-review is double-blind.

#### Processing Data citations

When data citations are present in the reference list of the manuscript, these should be processed in the same way as other references by the publisher. This means that formatting and quality control should take place at the production stage (see JATS4R data citation recommendations^26^).

#### DOIs and Compact Identifiers

Digital Object Identifiers (DOIs) are well understood by publishers as identifiers. DOIs are also assigned by many repositories to identify datasets. When available, they should be included in the data reference similarly to the use of DOIs for article references. An advantage to DOIs for data is that the associated metadata is centrally managed by the DataCite organization, similarly to how Crossref manages article metadata. DataCite and Crossref collaborate closely.

However, many domain-specific repositories in biomedical research do not issue DOIs, instead they issue locally-assigned identifiers (“accessions”, “accession numbers”). Funders of biomedical research may require data to be deposited in domain specific repositories e.g. GEO, dbGAP, and SRA, many of which use such locally resolvable accession numbers in lieu of DOIs.

Prior informal practice had been to qualify these by a leading prefix, so that the identifier became unique. In 2012 the European Molecular Biology Laboratory-European Bioinformatics Institute (EMBL-EBI) began tracking and issuing formal namespace prefixes to avoid collisions and support formal resolution on the Web ^27^. Subsequent efforts developed a collaborative curation model ^28^.

Now EMBL-EBI and the California Digital Library (CDL) maintain a common shared namespace registry and resolvers capable of interpreting and resolving PREFIX:ACCESSION patterns, or “compact identifiers”, hosted at these leading institutions ^14^. Technical work to develop the common repository and resolution rules was coordinated with the work of our Publishers Roadmap team.

This means that compact identifiers have now been formalized, are institutionally supported in the U.S. and in Europe, and may be used by in place of DOIs. We recommend this be done (1) where the repository does not issue DOIs for deposited datasets and (2) where the repository’s prefix has been registered. Similar to DOIs, compact identifiers are dereferenced by resolvers hosted at well-known resolver web addresses: http://identifiers.org (EMBL-EBI) and http://n2t.net (CDL). These resolver addresses, for example, both resolve the Gene Expression Omnibus local accession number GDS5157 (as https://identifiers.org/GEO:GDS5157 or https://n2t.net/GEO:GDS5157) to a primary expression dataset generated on the Illumina MouseWG-6 v2.0 expression beadchip, supporting findings on genetics of fear expression in an article by Andero et al. 2014 ^29^).

While these resources are still under active development to resolve an increasing number of identifiers, ensuring that either a DOI or a Compact Identifier is associated with data references will be important to support automatic resolution of these identifiers by software tools, which benefits authors, data providers and service providers. Other working group efforts are underway within the Research Data Alliance (RDA), for example in the Scholarly Link Exchange (Scholix) project (https://www.rd-alliance.org/groups/rdawds-scholarly-link-exchange-scholix-wg); and in other efforts such as THOR and FREYA (funded by the European Commission) to ensure the infrastructure to enable accurate and expedient resolvable linking between publishers, referenced datasets, and repositories.

### 3. Production

The main relevant components of the production process are the input from the peer review process (typically author manuscript in Word or LaTex files), conversion of this to XML and other formats (such as PDF, ePub), and the author proofing stage. Following all the preceding recommendations for the editorial process, the production process needs to identify relevant content and convert it to XML.

#### Data citations

The production department and its vendor(s) must ensure all data citations provided by the author in the reference list are processed appropriately using the correct XML tags. Typesetters must be provided with detailed instructions to achieve this. It is out of the scope of this paper to provide tools to identify datasets that are alluded to in a manuscript but are not present in the reference list; however, simple search and find commands could be executed using common terms and common database names.

##### XML requirements of data citations

For publishers using NISO standard JATS, version 1.1 and upwards, the JATS4R recommendation on data citations should be followed. The main other publisher-specific DTDs contain similar elements to allow for correct XML tagging.

Examples:

- eLIFE recommendation: https://github.com/elifesciences/XML-mapping/blob/master/elife-00666.xml
- JATS4R recommendation and examples: http://jats4r.org/data-citations

#### Data availability statement (DAS)

Output format from the editorial process will inform the production department as to how to identify and handle this content. For instance, some publishers require authors to provide the details within the submission screens and thus can output structured data from the submission system to production, others require a separate Word file to be uploaded, and others request the authors include this information in the manuscript file. Depending on the method used, production will need to process and convert this content accordingly.

Where the DAS will be contained/displayed within the PDF/ePub format of the article is decided by the individual publisher and this group will not provide recommendations for this.

### 4. Publication

#### Display Data Citations in the article

There are two primary methods of displaying data citations in a manuscript--in a separate data citations section or in the main references section. A separate data citations section promotes visibility, but inclusion in the main references section helps establish equal standing between data citations and standard references, and aids machine readable recovery, so is recommended.

Data citations should include a PID (such as a DOI) and should ideally include the minimum information recommended by DataCite and the FORCE11 data citation principles (Author, year, title, PID, repository). Where possible, PIDs should be favored over URLs, and they should function as links that resolve to the landing page of the dataset. Optionally, some publishers may choose to highlight the datasets on which the study relies by visualizing these.

#### Data Availability Statements

Data Availability Statements (DAS) should be rendered in the article (see Figure 3).

#### Downstream delivery to Crossref

Crossref ensures that links to scholarly literature persist over time through the Digital Object Identifier (DOI). They also provide infrastructure to the community by linking the publications to associated works and resources through the metadata that publishers deposit at publication, making research easy to find, cite, link, and assess. Links to data resources (i.e., data citations) are a core part of this service.

Publishers deposit the data citations by adding them directly into the standard metadata deposit via references and/or relation type. This is part of the existing content registration process and requires no new workflows. In the metadata deposit, publishers can deposit the literature-data links in two places: bibliographic references and/or relationships component. Since each has its own distinct advantage, both are encouraged where possible (see the Data Citations Deposit Guide linked below).

1. Bibliographic references: Publishers include the data citation into the deposit of bibliographic references for each publication. Here, publishers follow the general process for depositing references and apply tags to structure the metadata as applicable.
2. Relation type: Publishers assert the data citation in an existing section of the metadata deposit dedicated to connecting the publication to a variety of research objects associated with it (e.g., data and software, supporting information, protocols, videos, published peer reviews, preprint, conference papers, etc.). In addition to structured metadata about the data, this also allows publishers to identify any data linked as a direct output of the research results (viz., for scientific validation) if this is known.

When all of this information is sent to Crossref, extended opportunities for data mining and building up pictures of data citations, linking, and relationships will be possible. This also means that information is open and links between articles and datasets become available to other services that increase the discoverability of data and potential for reuse. For example, by sending these data citations to CrossRef, they become available in a Scholix compliant way (www.scholix.org) which enables their retrieval through the Data Literature Interlinking service (https://dliservice.research-infrastructures.eu/#/) - an easy way for both publishers and repositories to retrieve information about associations between articles and datasets.

More detail can be found in the Data & Software Citations Deposit Guide^30^.

#### Downstream delivery to PubMed

Metadata about data linked as a direct output of the research results can be deposited with the PubMed record for a research article for inclusion within PubMed, which maintains a controlled list of allowed databases here:

https://www.ncbi.nlm.nih.gov/books/NBK3828/#publisherhelp.Object_O Here is a tagging example:

<Object Type="Dryad">

<Param Name="id">10.5061/dryad.2f050</Param>

</Object>

#### Next steps

Several publishers are now in the process of implementing the JDDCP in line with the steps described in this roadmap. More work is still needed, both by individual publishers and by this group. This document describes basic steps that should be taken to enable authors to cite datasets. As a next step, improved workflows and tools should be developed to automate data citation further. In addition, authors need to be made aware of the importance of data citation and will require guidance on how to cite data. Ongoing coordination amongst publishers, data repositories, and other important stakeholders will be essential to ensure data is recognized as a primary research output. Table 2 outlines the implementation timelines of the different publishers that participated in this project.

**Table 2:** Estimated data citation implementation timelines for eight academic publishers. NB. All relevant journals at a given publisher will be included in the data citation rollout.

| **Publisher** | **Planning** | **Implementation** | **Fine-tuning** |
| --- | --- | --- | --- |
| eLife | Q1-Q3 2017 | Q4 2017 | Q2 2018 |
| Elsevier | Q2-Q3 2016 | Q4 2016 | 2017 |
| EMBO Press | Q1-Q2 2017 | Q3 2017 - Q1 2018 | Q1 2018 |
| Frontiers | Q1 - Q4 2017 | Q1 - Q2 2018 | 2018 |
| PLOS | Q1 - Q4 2017 | Q1 2018 | 2018 |
| SpringerNature | 2016 | Q4 2017 | Q1 2018 |
| Taylor & Francis | Q1-Q2 2017 | Q4 2017 continuing through 2018 | 2018 |
| Wiley | Q1 - Q2 2017 | Q3, Q4 2017 continuing through 2018 | 2018 |

## Discussion

This roadmap originated through the implementation phase of a project aimed at enhancing the reproducibility of scientific research and increasing credit for and reuse of data through data citation. The project was organized as a series of Working Groups in FORCE11 (https://force11.org/), an international organization of researchers, funders, publishers, librarians, and others seeking to improve digital research communication and eScholarship.

The effort began with the Joint Declaration of Data Citation Principles ^7,31^, which distilled and harmonized conclusions of significant prior studies by science policy bodies on how research data should be made available in digital scholarly communications. In the implementation phase (the Data Citation Implementation Pilot, (https://www.force11.org/group/dcip), repositories, publishers, and data centers formed three Expert Groups, respectively, with the aim of creating clear recommendations for implementing data citation in line with the JDDCP.

Once the steps outlined in this roadmap are implemented, authors will be able to cite datasets in the same way as they cite articles. In addition to ‘author’, ‘year’, and ‘title’, they will need to add the data repository, version and persistent unique identifier to ensure other researchers can unambiguously identify datasets. Publishers will be able to recognize the references as data references and process these accordingly, so that it becomes possible for data citations to be counted and for researchers to get credit for their work. These are essential steps for substantially increasing the FAIRness^6^ of research data. We believe this will in turn lead to better, more reproducible, and re-usable science and scholarship, with many benefits to society.

## Methods

In a series of teleconferences over a period of a year, major publishers compared current workflows and processes around data citation. Challenges were identified and recommendations structured according to the publisher workflows were drafted. In July 2016 this group met with additional representatives from publishers, researchers, funders, and not-for-profit open science organizations in order to resolve remaining challenges, validate recommendations, and to identify future tasks for development. From this the first full draft of the Publisher Roadmap was created. Feedback was then solicited and incorporated from other relevant stakeholders in the community as well as the other Data Citation Implementation Pilot working groups.

## Author contributions

*Helena Cousijn* and *Amye Kenall* co-chaired the DCIP Publishers Expert Group which produced this article. They had primary responsibility for leading regular telecons as well as a face-to-face meeting of participants (see Acknowledgements) at the SpringerNature London campus in July of 2016. Dr. Cousijn and Ms. Kenall provided the article structure; organized their Expert Group to collect and integrate information from the participating publishers, including their own organizations; and did the majority of writing for this article. They made equal contributions to the work.

*Emma Ganley, Patrick Polischuk, Melissa Harrison, David Kernohan, Thomas Lemberger, Simone Taylor* and *Fiona Murphy* participated in the work of the Publishers Expert Group and co-authored this article. They provided knowledgeable content and input to the work from the perspectives of their respective organizations. In addition, *Melissa Harrison* coordinated and informed this work with the perspective of the JATS4R group (Journal Article Tag Suite for Reuse), which she chairs.

*Tim Clark* coordinated the work of the Publishers Expert Group with the other DCIP participants (Repositories, Identifiers, JATS, and Primer/FAQ), co-authored sections of this article and edited the whole.

*Tim Clark* and *Maryann Martone* co-led the Data Citation Implementation Pilot as a whole.

## Acknowledgments

Research reported in this publication was supported in part by the National Institutes of Health under award number U24HL126127. The content is solely the responsibility of the authors and does not necessarily represent the official views of the National Institutes of Health.

The authors gratefully acknowledge the following members of the FORCE11/bioCADDIE Data Citation Pilot Publishers Expert Group, who participated in workshops and/or telecons to assist in developing this Roadmap: Helen Atkins (Public Library of Science); Paul Donohoe (SpringerNature); Scott Edmunds (GigaScience); Martin Fenner (DataCite); Ian Fore (National Cancer Institute, National Institutes of Health); Carole Goble (University of Manchester); Florian Graef (European Bioinformatics Institute); Iain Hrynaszkiewicz (SpringerNature); Johanna McEntyre (European Bioinformatics Institute); Ashlynne Merrifield (Taylor and Francis); Eleonora Presani (Elsevier); Perpetua Socorro (Frontiers); Caroline Sutton (Taylor and Francis); Michael Taylor (Digital Science).

## References

1. Uhlir, P. (ed.) For attribution: developing data attribution and citation practices and standards: summary of an international workshop. (National Academies, Washington DC, 2012).

2. CODATA/ITSCI Task Force on Data Citation. Out of cite, out of mind: the current state of practice, policy and technology for data citation. Data Science Journal 12, 1–75, doi: 10.2481/dsj.OSOM13-043 (2013).

3. Hodson, S. & Molly, L. Current best practice for research data management policies. Zenodo, doi:10.5281/zenodo.27872 (2015).

4. Committee on Ensuring the Utility and Integrity of Research Data in a Digital Age. Ensuring the integrity, accessibility, and stewardship of research data in the digital age. (The National Academies Press, 2009).

5. Royal Society. Science as an open enterprise. (The Royal Society Science Policy Center, London, 2012).

6. Wilkinson, M. D. et al. The FAIR Guiding Principles for scientific data management and stewardship. Scientific Data 3, 160018, doi:10.1038/sdata.2016.18 (2016).

7. Data Citation Synthesis Group. Joint declaration of data citation principles. FORCE11, doi:10.25490/a97f-egyk (2014).

8. Starr, J. et al. Achieving human and machine accessibility of cited data in scholarly publications. PeerJ 1:e1, doi:10.7717/peerj-cs.1 (2015).

9. Bierer, B. E., Crosas, M. & Pierce, H. H. Data authorship as an incentive to data sharing. The New England journal of medicine 377, 402, doi:10.1056/NEJMc1707245 (2017).

10. Vocile, B. Open science trends you need to know about, in Discover the Future of Research <(The Wiley Network, 2017).

11. Michener, W. K. Ecological data sharing. Ecological Informatics 29, 33–44, doi:10.1016/j.ecoinf.2015.06.010 (2015).

12. Piwowar, H. A., Day, R. S. & Fridsma, D. B. Sharing detailed research data is associated with increased citation rate. PLOS ONE 2, e308, doi:10.1371/journal.pone.0000308 (2007).

13. McKiernan, E. C. et al. How open science helps researchers succeed. eLife 5, e16800, doi:10.7554/eLife.16800,(2016).

14. Wimalaratne, S. M. et al. Uniform resolution of compact identifiers for biomedical data. Scientific Data [in press, this issue] (2018).

15. Fenner, M. et al. A data citation roadmap for scholarly data repositories. bioRxiv 097196, doi: 10.1101/097196 (2017).

16. McNutt, M. et al. Transparency in authors’ contributions and responsibilities to promote integrity in scientific publication. bioRxiv 140228, doi: 10.1101/140228 (2017).

17. Wiley. Sharing and citing your research data, <https://authorservices.wiley.com/author-resources/Journal-Authors/licensing-open-access/open-access/data-sharing.html> (2018).

18. Springer Nature. Research data policy types, <http://www.springernature.com/gp/group/data-policy/policy-types> (2018).

19. Elsevier. Research data guidelines, <https://www.elsevier.com/authors/author-services/research-data/data-guidelines> (2018).

20. Coalition on Publishing Data in the Earth and Space Sciences. COPDESS suggested author instructions and best practices for journals. <http://www.copdess.org/copdess-suggested-author-instructions-and-best-practices-for-journals/> (accessed Feb 11, 2018).

21. Bloom, T., Ganley, E. & Winker, M. Data access for the open access literature: PLOS’s data policy. PLOS Biology 12, e1001797, doi:10.1371/journal.pbio.1001797 (2014).

22. Kim, K. W. et al. Coordinated inhibition of C/EBP by Tribbles in multiple tissues is essential for Caenorhabditis elegans development. BMC Biology 14, 104, doi:10.1186/s12915-016-0320-z (2016). 1

23. Edgar, R., Domrachev, M. & Lash, A. Gene expression omnibus: NCBI gene expression and hybridization array data repository. Nucleic Acids Res 30, 207 –210 (2002).

24. Barrett, T. et al. NCBI GEO: archive for functional genomics data sets—10 years on. Nucleic Acids Research 39, D1005–D1010, doi:10.1093/nar/gkq1184 (2011).

25. Sarkans, U. et al. The BioStudies database—one stop shop for all data supporting a life sciences study. Nucleic Acids Research 46, D1266–D1270, doi:10.1093/nar/gkx965 (2018).

26. JATS4R. Data citations, <https://jats4r.org/data-citations> (2018).

27. Juty, N., Le Novère, N. & Laibe, C. Identifiers.org and MIRIAM registry: community resources to provide persistent identification. Nucleic Acids Research 40, D580–D586, doi:10.1093/nar/gkr1097 (2012).

28. Juty, N., Le Novère, N., Hermjakob, H. & Laibe, C. Towards the collaborative Curation of the Registry underlying identifiers.org. Database 2013, bat017–bat017, doi:10.1093/database/bat017 (2013).

29. Andero, R., Dias, Brian G. & Ressler, Kerry J. A role for Tac2, NkB, and Nk3 receptor in normal and dysregulated fear memory consolidation. Neuron 83, 444–454, doi:10.1016/j.neuron.2014.05.028 (2014)

30. Crossref. Crossref data & software citation deposit guide for publishers, <https://support.crossref.org/hc/en-us/articles/215787303-Crossref-Data-Software-Citation-Deposit-Guide-for-Publishers> (2018).

31. Altman, M., Borgman, C., Crosas, M. & Martone, M. An introduction to the joint principles for data citation. Bulletin of the Association for Information Science and Technology 41, 43–45, doi:10.1002/bult.2015.1720410313 (2015).

